# *Pseudomonas syringae* addresses distinct environmental challenges during plant infection through the coordinated deployment of polysaccharides

**DOI:** 10.1101/2021.06.18.449010

**Authors:** Pilla Sankara Krishna, Stuart Daniel Woodcock, Sebastian Pfeilmeier, Stephen Bornemann, Cyril Zipfel, Jacob George Malone

**Affiliations:** Department of Molecular Microbiology, John Innes Centre, Norwich Research Park, Norwich, United Kingdom NR4 7UH; The Sainsbury Laboratory, University of East Anglia, Norwich Research Park, Norwich, United Kingdom NR4 7UH; Department of Biological Chemistry, John Innes Centre, Norwich Research Park, Norwich, United Kingdom NR4 7UH; University of East Anglia, Norwich Research Park, Norwich, United Kingdom NR4 7TJ

**Keywords:** Exopolysaccharides, *Pseudomonas syringae*, lipopolysaccharide, cyclic di-GMP, biofilm, surface adhesion, plant infection

## Abstract

Prior to infection, phytopathogenic bacteria face a challenging environment on the plant surface, where they are exposed to nutrient starvation and abiotic stresses. Pathways enabling surface adhesion, stress tolerance and epiphytic survival are important for successful plant pathogenesis. Understanding the roles and regulation of these pathways is therefore crucial to fully understand bacterial plant infections. The phytopathogen *Pseudomonas syringae* pv. *tomato* (*Pst*) encodes multiple polysaccharides that are implicated in biofilm formation, stress survival and virulence in other microbes. To examine how these polysaccharides impact *Pst* epiphytic survival and pathogenesis, we analysed mutants in multiple polysaccharide loci to determine their intersecting contributions to epiphytic survival and infection. In parallel, we used qRT-PCR to analyse the regulation of each pathway. *Pst* polysaccharides are tightly coordinated by multiple environmental signals. Nutrient availability, temperature and surface association strongly affect the expression of different polysaccharides under the control of the signalling proteins *ladS* and *cbrB* and the second messenger cyclic-di-GMP. Furthermore, functionally redundant, combinatorial phenotypes were observed for several polysaccharides. Exopolysaccharides and WapQ-mediated lipopolysaccharide production are important for leaf adhesion, while α-glucan and alginate together confer desiccation tolerance. Our results suggest that polysaccharides play important roles in overcoming environmental challenges to *Pst* during plant infection.

**Highlight:** *Pseudomonas syringae* uses the coordinated deployment of polysaccharides to address environmental challenges during plant colonization. Functional redundancy renders individual polysaccharides dispensable during plant infection, but their combined loss impedes pathogenicity.

## Introduction

*Pseudomonas syringae* is a Gram negative, plant pathogenic bacterium and a widely used model system to understand plant-microbe interactions (Hirano and Upper, 2000; Xin *et al*., 2018). Taxonomic studies indicate that *P. syringae* is in fact a diverse phylogenetic group containing more than 15 species and over 60 pathovars (Young, 2010). Pathovars of *P. syringae* infect almost all economically important crop plants and are considered to be one of the most common and damaging bacterial plant pathogens that infect the phyllosphere (Xin *et al*., 2018). *P. syringae* pathogenesis is attributed to a repertoire of virulence factors, including secretion systems, phytotoxins and phytohormones, quorum sensing pathways, ice nucleation agents, cell wall-degrading enzymes and exopolysaccharides (EPS) (Hirano and Upper, 2000; Pfeilmeier *et al*., 2016a; Xin *et al*., 2018). Plant infection by *P. syringae* consists of epiphytic and endophytic phases (Xin and He, 2013). Survival on plant surfaces like leaves, stems or fruits is referred as the epiphytic phase, while the endophytic phase describes bacterial entry into the plant tissue and colonization of the intercellular apoplastic space. Bacteria face a challenging environment on the plant surface, where they are routinely exposed to mechanical, temperature and desiccation stresses, nutrient starvation, and ultraviolet irradiation (Andrews and Harris, 2000; Djonovic *et al*., 2013; Pfeilmeier *et al*., 2016a; Wu *et al*., 2012). Thus, adhesion to plant surfaces, stress tolerance and epiphytic survival are likely to be important for successful pathogenesis.

Extracellular matrix and cell envelope components such as lipopolysaccharide (LPS) and EPS molecules interact directly with the host/plant surface and hence are vital in establishing an interaction during the epiphytic phase (King and Roberts, 2016; Mhedbi-Hajri *et al*., 2011). Genetic studies have shown that lipopolysaccharide production is required for efficient host colonization and for full virulence in plant pathogenic *Pseudomonas* spp., *Erwinia amylovora* and *Burkholderia cenocepacia* (Berry *et al*., 2009; Khodai-Kalaki *et al*., 2015; Kutschera *et al*., 2019). Similarly, EPS production has been implicated in epiphytic survival of *P. syringae* and *Xanthomonas* spp., and in the wider plant microbiome (Dunger *et al*., 2007; Yu *et al*., 1999). It has also been suggested to contribute to bacterial freeze–thaw resistance (Wu *et al*., 2012). The widely studied model plant pathogen *P. syringae* pv. *tomato* str. DC3000 (*Pst*) contains multiple polysaccharide gene clusters that can potentially contribute to biofilm matrix formation including alginate, Wss cellulose, Psl and α-glucan (Winsor *et al*., 2016).

Alginate is a copolymer of acetylated β-1,4-linked D-mannuronic acid and L-glucuronic acid that contributes to antibiotic and immune system protection in the human pathogen *P. aeruginosa* (Hentzer *et al*., 2001; Pier *et al*., 2001). Alginate is not considered essential to *Pseudomonas* biofilm formation (Laue *et al*., 2006; Wozniak *et al*., 2003), but has been implicated in epiphytic fitness and virulence in some *P. syringae* strains (Helmann *et al*., 2019; McAtee *et al*., 2018; Yu *et al*., 1999). Cellulose is a homopolymer made of β-D-glucose monomers and is a primary component of the biofilm matrix of *Pst* and many other bacteria (Farias *et al*., 2019; Prada-Ramirez *et al*., 2016; Serra *et al*., 2013). Cellulose has been suggested to play an important role in the transition between epiphytic and pathogenic phases of leaf association (Arrebola *et al*., 2015). Similarly, acetylated cellulose (Wss) contributes substantially to root association of the rhizosphere bacterium *P. fluorescens* (Gal *et al*., 2003). Psl is a pentasaccharide polymer of D-mannose, D-glucose and L-rhamnose subunits that plays essential roles in biofilm formation, adhesion, motility, and stress protection in *P. aeruginosa* (Billings *et al*., 2013; Byrd *et al*., 2009; Periasamy *et al*., 2015). The function of Psl in *P. syringae* remains relatively unclear, although it has recently been implicated in swarming motility and pathogenicity regulation in the *P. syringae* mango pathovar UMAF0158 (Heredia-Ponce *et al*., 2020).

GlgE derived α-glucan is a glycogen-like homopolymer of glucose monomers with α-1,4 glycosidic links and α-1,6 linked branch points (Edstrom, 1972). α-Glucan is a ubiquitous carbon store and plays important roles in desiccation stress tolerance in *P. aeruginosa* (Woodcock *et al*., 2021), and virulence in *Mycobacterium tuberculosis* (Koliwer-Brandl *et al*., 2016; Sambou *et al*., 2008). Recently, α-glucan has been implicated as an EPS in the virulent kiwi pathovar *P. actinidiae* (Ghods *et al*., 2015). The ubiquitous compatible solute trehalose is not only a precursor of α-glucan (Woodcock *et al*., 2021) but also contributes to osmotic stress tolerance during epiphytic survival by *P. syringae* (Freeman *et al*., 2010). Finally, levan is a β-2,6 polyfructan with extensive branching through β-2,1 linkages (Laue *et al*., 2006; Li and Ullrich, 2001). Levan has not been implicated in biofilm formation or epiphytic survival. Instead, it has been suggested to function as a storage molecule that may be produced in the apoplast (Laue *et al*., 2006; Yu *et al*., 2013).

LPS molecules are complex glycoconjugate molecules whose biosynthesis, packing and role in pathogenesis and immune evasion is well understood in animal/human pathosystems but less explored in plant pathogens. The *Pst* genome encodes at least four lipopolysaccharide biosynthesis operons, including genes for Waap and Waa proteins alongside *wapQ/wapG* and *wbpL* (Winsor *et al*., 2016). In *Erwinia amylovora*, LPS production contributes to virulence and oxidative stress protection (Berry *et al*., 2009). Similarly, in *Burkholderia cenocepacia* and *Xylella fastidiosa* virulence is compromised in LPS mutants (Khodai-Kalaki *et al*., 2015; Rapicavoli *et al*., 2018). Recent studies have shown that *Pst* cells lacking a *wbpL* orthologue failed to produce O-polysaccharide and exhibited reduced apoplast colonization and pathogenesis (Kutschera *et al*., 2019). Conversely, the LPS core kinase gene *waaP* can be deleted in several phytopathogenic *Pseudomonas* strains but not in *P. aeruginosa* (DeLucia *et al*., 2011; Kutschera *et al*., 2019). Finally, *wapQ* encodes an LPS kinase but is poorly characterised in *P. syringae*.

*Pst* phytotoxins and effector proteins and their roles in immune suppression have been extensively studied. However, to obtain a comprehensive understanding of plant infection, we also need to unravel the relationship between bacterial survival, stress tolerance and pathogenesis. In this study, we build on prior research on cellulose and alginate production in *Pst* (Farias *et al*., 2019; Keith *et al*., 2003; Perez-Mendoza *et al*., 2019) to examine the intersecting roles of multiple *Pst* polysaccharide molecules in mediating bacterial infection and enabling epiphytic survival. *Pst* mutants in the alginate (*alg*), *psl*, cellulose (*wss*), α-glucan (*glg/tre*) and *wapQ* pathways were constructed, and their relative contributions to phenotypes including colony morphology, abiotic stress tolerance, epiphytic survival, and plant infection were defined.

The polysaccharide pathways in *Pst* are each tightly regulated by external environmental cues. In particular, *alg* and *psl* expression were shown to be strongly nutrient dependent, while *wss* production was activated by low temperatures and overproduction of the second messenger molecule cyclic-di-GMP (cdG) (Jenal *et al*., 2017). Expression of the *glg* trehalose/α-glucan locus is induced by surface association, under the control of the global regulators *ladS* and *cbrB* (Grenga *et al*., 2017; Sonnleitner *et al*., 2009). Strikingly, we observed a substantial degree of functional redundancy and phenotypic interaction between the polysaccharide molecules in our study. While disruption of individual polysaccharide loci had little effect on *Pst* plant infection, mutation of multiple loci led to compromised infection or delayed disease onset for several different combinations. Similarly, we observed combinatorial phenotypic effects for several loci, suggestive of functional redundancy. Alginate and α-glucan combine to confer desiccation stress tolerance, while EPS and WapQ mediated LPS production together contribute to leaf surface adhesion. Our data suggest that bacterial polysaccharides play important, intersecting roles in enabling infection by plant pathogens.

## Methods

### Strains and growth conditions

Unless otherwise stated all strains were grown at 28 °C with shaking. Bacterial strains and plasmids used in this study are listed in Table 3. King’s B medium (KB) (King *et al*., 1954), L medium (Sambrook and Russel, 2001), soya flour mannitol Medium (SFM) (2% soya flour and 2% mannitol) and M9 medium (M9 salts (Sambrook and Russel, 2001) supplemented with 0.4% glucose, 0.4% casamino acids and 50 µM FeCl_3_) were used for culturing and for *in vitro* assays as described in the text. Antibiotics were used at a final concentration of gentamicin at 25 μg ml^-1^, tetracycline at 12.5 μg ml^-1^and kanamycin 25 μg ml^-1^ for selection of mutants or during genetic manipulations. Final concentration of X-gal 40 μg ml^-1^ and IPTG 1mM were used for blue-white screening of reporter strain.

### Mutagenesis and genetic manipulation

Gene deletion vectors were constructed by amplifying the upstream and downstream regions flanking the desired gene from the *Pst* genome using primers No. 1-72, listed in Table 4. Amplified up and downstream flanking regions were then cloned into the multiple cloning site of the suicide vector pTS1 (Scott *et al*., 2017). *Pst* cells were transformed using electroporation with the appropriate deletion vectors following the method described in (Choi *et al*., 2006). Single crossover integrations of each plasmid into the chromosome were selected on tetracycline plates and re-streaked before single colonies were grown in overnight in KB media without selection. Double crossovers were counter-selected by plating serial dilutions onto L agar containing 10% (w/v) sucrose. Deletion mutants were confirmed by PCR using corresponding primers labelled test-F and test-R in (Table 4). To produce the *Pst*-P*glgA-lacZ* reporter, the promoter region of *Pst glgA* (*PSPTO3125*) including several codons of the open reading frame was amplified using primers No.73-74, listed in Table 4 and cloned between the BamHI and HindIII sites of pUC18-mini-*Tn7*T-Gm-LacZ10 (Choi and Schweizer, 2006).

### Transposon mutagenesis screening

The plasmid pALMAR3 was introduced into *Pst*-P*glgA-lacZ* via biparental mating with *E. coli* S17-1. Mariner transposon insertion mutants were selected by plating onto L agar containing gentamycin, tetracycline and X-Gal. Colonies showing changes in LacZ activity were re-streaked and the location of the transposon in each case was determined by arbitrary PCR (O’Toole and Kolter, 1998) using primers (75-78) listed in Table 4.

### Extraction and NMR analysis of metabolites

Overnight *Pst* cultures were grown in M9 medium, then adjusted to a cell density of 0.5 (OD_600_) in PBS. Mixed cellulose ester filter discs (Merck Millipore) were placed on the surface of M9 agar plates and coated with the diluted cell suspensions. Plates were then incubated at 28 °C for 48 hours before cells were harvested from each disc and resuspended in 5 ml of ddH_2_O in a 15 ml plastic tube by vigorous vortexing. Vacuum dried cells were weighed (dry weight), resuspended in ddH_2_O and boiled at 95 °C for 20 minutes. Boiled cells were then centrifuged (10,000 × g, 10 minutes) and the resulting supernatant was subject to additional boiling and centrifugation steps to remove any remaining insoluble contaminants. The supernatant, now containing the total water-soluble metabolite content of the cell sample, was dried under vacuum for 16 hours. The resulting dried pellet was resuspended in 1200 μl D_2_O and analysed by H^1^-NMR spectroscopy (Bruker AVANCE III 400 spectrometer), at room temperature with water suppression. Metabolite chemical shifts were recorded as parts per million (ppm) relative to 0.5 mM trimethylsilyl propanoic acid (TMSP; 0.00 ppm). Spectra were analysed using Topspin 3.0 (Bruker) and the concentrations of metabolites were established through the manual integration of peaks relative to TMSP. The resonances were assigned based on previously established spectra (Usui *et al*., 1974; Woodcock *et al*., 2021)

### Plant infection assays and bacterial load estimation

*Arabidopsis thaliana* Col-0 (Col0) plants were grown in controlled environment room (CER) in short day conditions: 10 h light, 22°C, 70% relative humidity. Cell cultures were grown overnight in L, then resuspended in 10mM MgCl_2_ and adjusted to a final density (OD_600_) of 0.2, equivalent to 1 × 10^8^ CFU ml^-1^. 0.04% (w/v) of Silwet® L-77 (phytotech labs) was added to the cell suspension as a surfactant just before spraying on the plants. Four-to five-week-old plants were infected using a hand-held sprayer until all leaves appeared wet. For leaf infiltration experiments, cells at a final OD_600_ of 0.02 (equivalent to 10^7^ cells ml^-1^) were gently infiltrated into the leaf apoplast using a 1 ml syringe until leaves appeared wet. For calculating bacterial load, two 7mm diameter leaf discs (area 0.384 cm^2^) were collected for each sample and decanted into 200 µl of sterile 10 mM MgCl_2._ Samples were lysed using a Geno/Grinder 2010 high throughput tissue homogenizer, with two 3mm glass beads and 2 cycles of 1700 vibrationsmin^-1^ with a 1 min break. Bacterial counts were determined by plating 10-fold serial dilutions on L agar plates with 50 µg ml^-1^ Rifampicin and 20 µg ml^-1^ Nystatin. A minimum of eight plants were used for each condition and assays were run at least twice independently. Bacterial load was calculated and presented as CFU per unit leaf area (CFUcm^-2^).

### Growth assays

Bacterial growth was measured in a microplate spectrometer using a minimum of 5 biological replicates and is presented as mean +/-standard deviation. 150 μl of the indicated growth medium in each case was added to the wells of clear-bottomed, black-walled 96-well microplates. Growth was initiated by the addition of 5 μl of overnight cell culture (L media), to obtain a starting OD_600_ of 0.01. Plates were incubated statically at 28 °C and agitated (500 RPM, 5 s) prior to each data acquisition step. Optical density was measured at 600 nm. Experiments were conducted at least twice independently.

### Osmotic stress assays

Overnight *Pst* cultures were grown in M9 medium, then diluted to OD_600_ of 0.01 in M9 medium. Five µl of each diluted sample was used to inoculate 150 µl of M9 medium. To examine osmotic stress conditions, growth media were supplemented with 0.35 M NaCl. Growth was monitored by measuring the optical density at 600 nm every hour for 6 days. Assays were conducted in triplicate and repeated at least twice independently, with a representative sample shown in each case.

### Desiccation tolerance assays

Desiccation tolerance assays were conducted following the method as previously described (Woodcock *et al*., 2021). Overnight *Pst* cultures were grown in M9 medium, then diluted to an OD_600_ of 0.1 in PBS. Ten µl of each culture were spotted onto 15 mm grade 1 Whatman filter discs. After drying for 1 minute at room temperature, discs were placed onto M9 agar plates and incubated at 28 °C for 4 hours to enable bacteria to recover and begin dividing.

After incubation, the filter discs were subjected to controlled desiccation in tightly sealed bell chambers containing either water (100% RH) or a saturated solution of NaCl (75% RH) (de Goffau *et al*., 2009) for 2 hours. Bacteria were then recovered from filter discs in 3 ml of PBS and serially diluted before spreading onto L agar plates. CFU were determined for each strain, then Log_10_(CFU) values analysed by linear mixed modelling using restricted maximum likelihood (REML) following the methodology described in (Woodcock et al., 2021). Assays were conducted in triplicate and repeated at least twice independently.

### Assessment of colony morphology

To assess the colony morphology of the *Pst* mutant strains, overnight cultures were grown in L media, resuspended in 10 mM MgCl_2_ solution and final densities were adjusted to an OD_600_ of 0.1. Agar plates were prepared and allowed to dry for 45 min in a sterile flow chamber. KB and SFM (Kieser and John Innes Foundation., 2000) plates were prepared, with Congo Red (CR) dye added as appropriate (30 μg ml^-1^). Five μl of each culture was spotted onto the agar surface and allowed to dry in a sterile flow chamber. Plates were then incubated at 28 °C for the indicated period before photographing, with a representative image shown in each case. For temperature treatments, plates were incubated at 28 °C for 24 h to enable colony establishment before plates were transferred to the designated temperature for a further incubation period as stated in the text.

### Leaf surface adhesion and surface survival assays

Adhesion assays were performed according to Arrebola et al., (Arrebola *et al*., 2015) with minor modifications. Tomato (cultivar Moneymaker) leaf discs (1 cm diameter) were prepared and placed in 70% ethanol for 30 sec with gentle swirling, then washed 3 times with 200 ml of sterile water. These leaf discs were placed on water agar with adaxial surface facing upwards and allowed to air dry in a sterile flow hood. Overnight bacterial cultures were adjusted to 10^8^ CFU ml^-1^ in 10 mM MgCl_2_. Drops (10□μl) of each strain were then inoculated on the adaxial surface of the leaf discs and allowed to air dry. After 5□h, the leaf pieces were gently washed by placing each disc in 1□ml of sterile 0.85% NaCl solution and gently inverting 3 to 4 times to remove unattached cells. 3 washed leaf discs were placed in 1□ml of sterile 0.85% NaCl, vortexed for 30□s to release adhered cells, followed by serial dilution and plating onto L agar with rifampicin (50 µg ml^-1^) and nystatin (20 µg ml^-1^) plates for CFU counting. Three such replicates for each strain and at least two independent experiments were performed. For *in vitro* leaf surface survival assays, Tomato (cultivar Moneymaker) leaf discs (1 cm diameter) were prepared and inoculated with bacteria as above. At respective time points leaf discs were collected and placed in 1□ml of sterile 10 mM MgCl_2_ vigorously vortexed for 30□s to release cells and CFU were counted following serial dilution.

### RNA isolation and qRT-PCR

Total RNA was extracted from cells grown on KB agar or SFM agar plates for 24 h at 28 °C. For low temperature treatment, 24 h old cultures were shifted from 28 °C to 8 °C and allowed to grow for another 24 h before collection. Cells scraped from plates were resuspended in 1 ml of RNA later and pelleted by centrifugation. RNA was isolated from pelleted cells using column capture (Qiagen RNeasy Mini Kit) following the manufacturer’s instructions. Purified RNA was subjected to additional DNase treatment (Turbo DNase, Ambion). RNA integrity was verified by agarose gel electrophoresis and the absence of genomic DNA contamination was confirmed by a negative response to 16S rRNA gene amplification in PCR reactions using isolated RNA as template. cDNA was prepared from isolated RNA using Superscript II reverse transcriptase (Invitrogen) following the manufacturer’s instructions. Gene specific primers were designed using the IDT primer quest tool, then qRT PCR assays were conducted in a Bio-Rad CFX96 Touch RT-PCR machine using a SensiFAST− SYBR® No-ROX Kit, with the following settings: 3 min at 95°C and 50 cycles of 5 sec at 95°C, 10 sec at 62°C and 10 sec at 72°C followed by a melting curve. Two independent RNA extractions and three technical replicates per extraction were assessed. *16S rRNA* and *gyrA* genes were used as internal controls for calculation of relative gene expression. Gene specific primer sequences (No. 79-92) used in this study were listed in (Table 4).

## Results

### Deletion of individual *Pst* polysaccharide loci has little effect on *Arabidopsis* infection

The *Pst* genome contains well-conserved predicted gene clusters for production of the polysaccharides alginate, Wss, Psl, α-glucan and levan, as well as WapP/WaaP LPS biosynthetic clusters (Winsor *et al*., 2016), summarised in Fig. S1-S6. Levan has been suggested to function as an apoplastic storage molecule (Laue *et al*., 2006; Yu *et al*., 2013) and is not implicated in biofilm or survival on leaf surfaces. Therefore, it was not studied further here. To probe the relationship between the remaining polysaccharide pathways and plant pathogenicity, we produced non-polar deletion mutants in key biosynthetic genes from each operon. Deletions in *algX-G* (Δ*alg*), *wssB-C* (Δ*wss*) and *pslD*-*E* (Δ*psl*), alongside a triple mutant of all three clusters (Δ*EPS*) have been described previously (Lammertz *et al*., 2019). *Pst* has similar arrangement of trehalose and α-glucan biosynthetic genes to *P. aeruginosa* PA01 (Woodcock *et al*., 2021), and we predict that the pathway is likely to function similarly in both species (Fig. S5, S6 & S8). Consequently, we produced mutants of the *treS* (Δ*PSPTO2760-62*) & *treY/Z* (Δ*PSPTO3125-3130*) operons. Finally, we deleted the well conserved LPS kinase gene *wapQ* (*PSPTO4998*) (Fig. S1, S7).

To investigate the role of individual polysaccharides on plant infection, we conducted a series of infection experiments in *Arabidopsis thaliana* Col-0 (Col-0) plants. No significant differences were observed between any of the *Pst* single mutant strains for spray infections (Fig. 1A). Previous research has highlighted the importance of the EPS alginate in apoplastic, but not epiphytic, survival in *P. syringae* B728a (Helmann *et al*., 2019). This suggests that bypassing the initial stages of infection may uncover differences between our mutants. To test this, we conducted leaf infiltration assays with our three *Pst* EPS single mutants. However, while a slight increase in necrosis symptoms was observed for leaves infected with the Δ*psl* mutant (Fig. 1B), no significant difference in bacterial load versus WT was seen for any of the single EPS mutants tested (Fig. 1C).

**Fig. 1.**
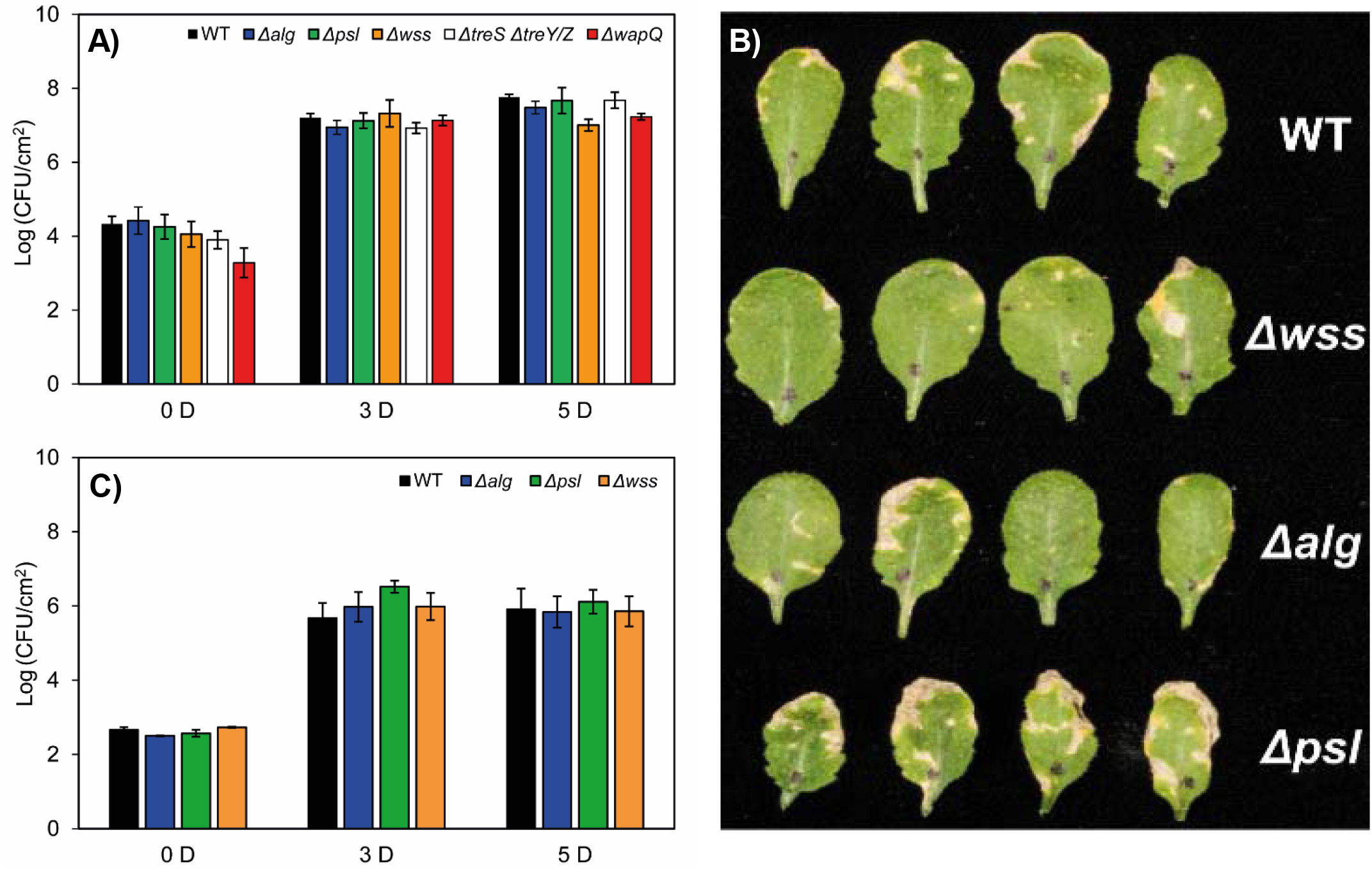
Deletion of individual polysaccharide loci has little effect on *Pst* Arabidopsis infections. (A) Bacterial load following spray infection at different days post infection [D]. (B) *Arabidopsis thaliana* Col0 leaves 5 days post-infiltration with different *Pst* mutants. (C) Bacterial load post-infection following infiltration at different days post infection [D]. Similar results were obtained for three biologically independent experiments in each case. Error bars represent mean ± SD from three technical replicates of a representative experiment.

### Wss, alginate, Psl and α-glucan production are regulated by cdG and nutrient availability

To understand why the *Pst* polysaccharide loci are apparently dispensable for plant infection, we tested their expression and production in response to a variety of intracellular and external signals. In general, disruption of individual polysaccharide loci did not lead to marked differences in *Pst* morphology for colonies grown on different nutrient media, with a few exceptions (Fig. 2A). On soy flour mannitol (SFM), the distinctive, alginate-dependent white mucoid colony phenotype was absent in the Δ*alg* mutant. Deletion of Δ*treS* Δ*treY/Z* produced noticeably thicker, more mucoid white colonies on SFM, suggesting a possible increased production of alginate in the absence of α-glucan. Deletion of *wapQ* produced a distinctive phenotype on KB Congo Red agar, with a dark halo surrounding the central zone of the *Pst* colony (Fig. 2A).

**Fig. 2.**
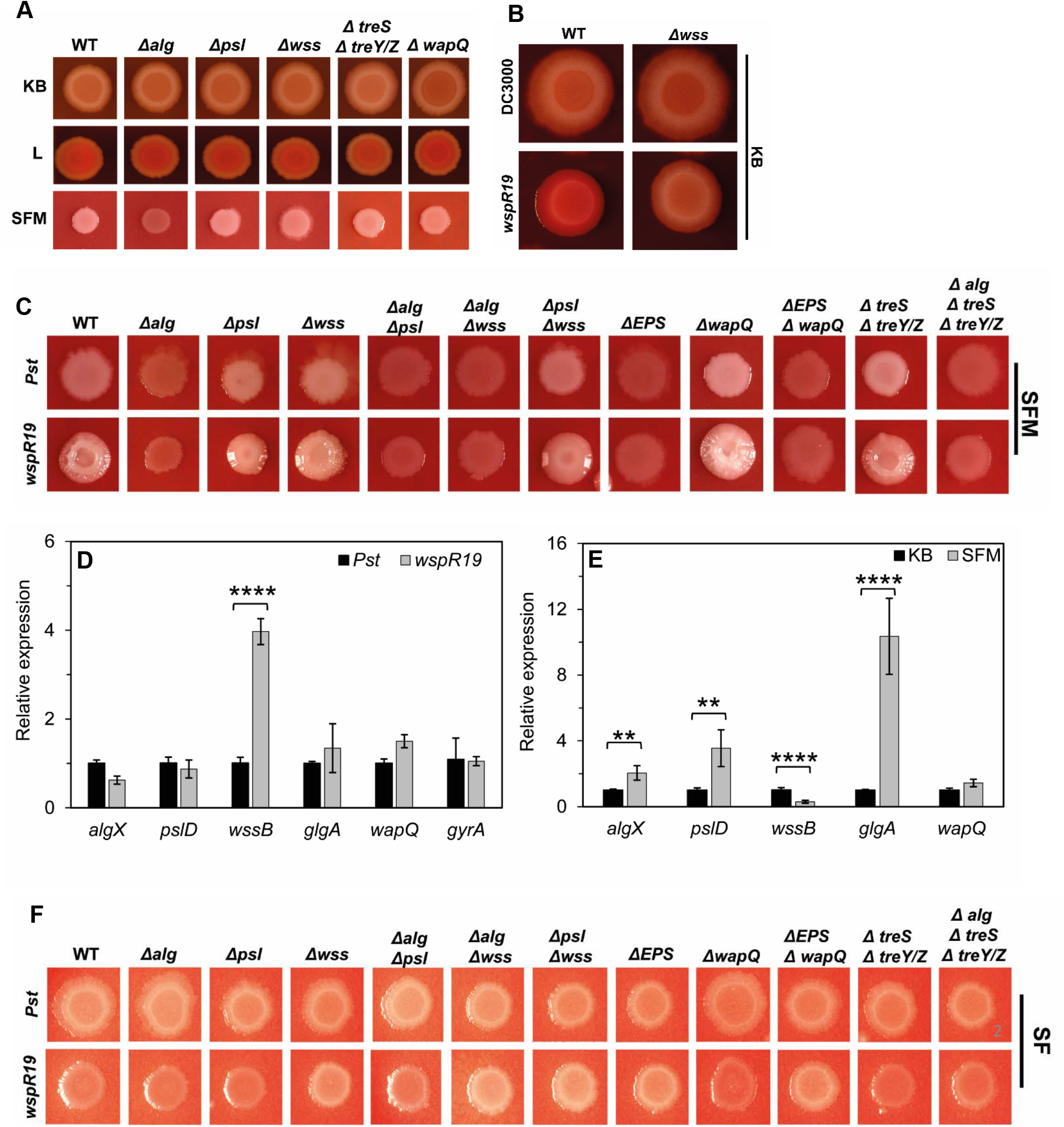
Wss, alginate, Psl and α-glucan production are stimulated by cdG and nutrient availability. A) Phenotypes of mutant strains on CR supplemented King’s B (KB), L medium (L) and soy flour mannitol (SFM) agar plates. (B) Effect of high cdG on *Pst* colony morphology at 28 °C on CR supplemented KB. (C) Colony morphology of *Pst* strains on CR supplemented soy flour mannitol medium (SFM). Strains with natural levels of cdG (*Pst*), strains with high c-di-GMP (*wspR19*). (D) Quantitative real time PCR analysis of the expression of polysaccharide producing genes in WT (*Pst*, black) and in the presence of high levels of cdG (*wspR19*, grey). (E) Quantitative real time PCR analysis of the expression of polysaccharide producing genes for WT *Pst* grown on KB agar (black) and SFM agar (grey). Error bars represent mean ± SD with n=2 (three technical replicates each) for (D) and (E). Significance as determined by student’s t-test (** indicates p<0.005 and **** indicates p<0.0001). (F) Colony morphology of *Pst* strains on CR supplemented soy flour (SF) medium. Similar results were obtained in two independent experiments and a representative picture is shown.

The second messenger cdG stimulates EPS production in many bacterial species including *Pst* (Jenal *et al*., 2017; Perez-Mendoza *et al*., 2019). To examine how cdG levels affect *Pst* polysaccharide production, we transformed our mutants with the plasmid pBBR-*wspR19*, which elevates cdG levels by approx. 15-fold in *Pst* (Pfeilmeier *et al*., 2016b). Curiously, unlike with many *Pseudomonas* strains (Malone *et al*., 2010) we saw little impact of cdG overproduction on colonies grown on KB agar at 28 °C, besides a slight increase in Congo Red (CR) binding that was absent in the Δ*wss* background (Fig. 2B, S9A). Conversely, growth on SFM led to the formation of mucoid, wrinkly colonies upon *wspR19* expression (Fig. 2C). Two polysaccharide gene deletions markedly affected this colony morphology. First, *wspR19+* Δ*psl* mutants produced smooth, mucoid colonies, supporting a role for Psl in maintaining the architecture of wrinkly colonies on SFM. Second, *alg* disruption abolished both mucoidy and the wrinkly phenotype in all backgrounds, implicating alginate as the primary structural EPS for *Pst* on SFM (Fig. 2C).

These results were supported by qRT-PCR of the *EPS* genes. *wssB* mRNA abundance increased markedly upon *wspR19* expression in *Pst* grown on KB agar, while the other tested genes were unaffected (Fig. 2D). However, a comparison of WT *Pst* grown on KB and SFM agar showed significantly higher *algX, pslD* and *glgA* mRNA abundance, and reduced levels of *wssB* mRNA for colonies on SFM, suggesting that expression of these EPS loci is strongly dependent on nutrient cues from the environment (Fig. 2E). The cdG-dependent, alginate-linked mucoid phenotype disappeared in the absence of mannitol, i.e., for strains grown on soy flour (SF) agar, suggesting that mannitol is a key nutrient for alginate production. Curiously, *Pst wspR19+* strains lacking the complete Wss, produced white slightly mucoid colonies on SF plates. This phenotype was independent of alginate, Psl or WapQ LPS production, suggesting that another, unknown polysaccharide may be upregulated under these conditions (Fig. 2F).

### Wss, Psl and α-glucan production are stimulated, while alginate is suppressed, by low temperature

Next, we examined the impact of temperature on polysaccharide gene expression and the associated changes in colony morphology. Switching the temperature from 28 °C to 8 °C produced relatively subtle changes in WT *Pst* but led to the formation of dry, wrinkled colonies upon *wspR19* expression on KB agar (Fig. 3A). The switch from 28 °C to 8 °C led to significant increases in mRNA abundance for *pslD* and *glgA* and reduced levels of *algX* (Fig. 3B). A similar pattern was seen for *Pst wspR19+*, with the exception of *wssB*, where a greater increase in mRNA abundance was seen at 8 °C than at 28 °C (compare Fig. 2D, 3B & 3D).

**Fig. 3.**
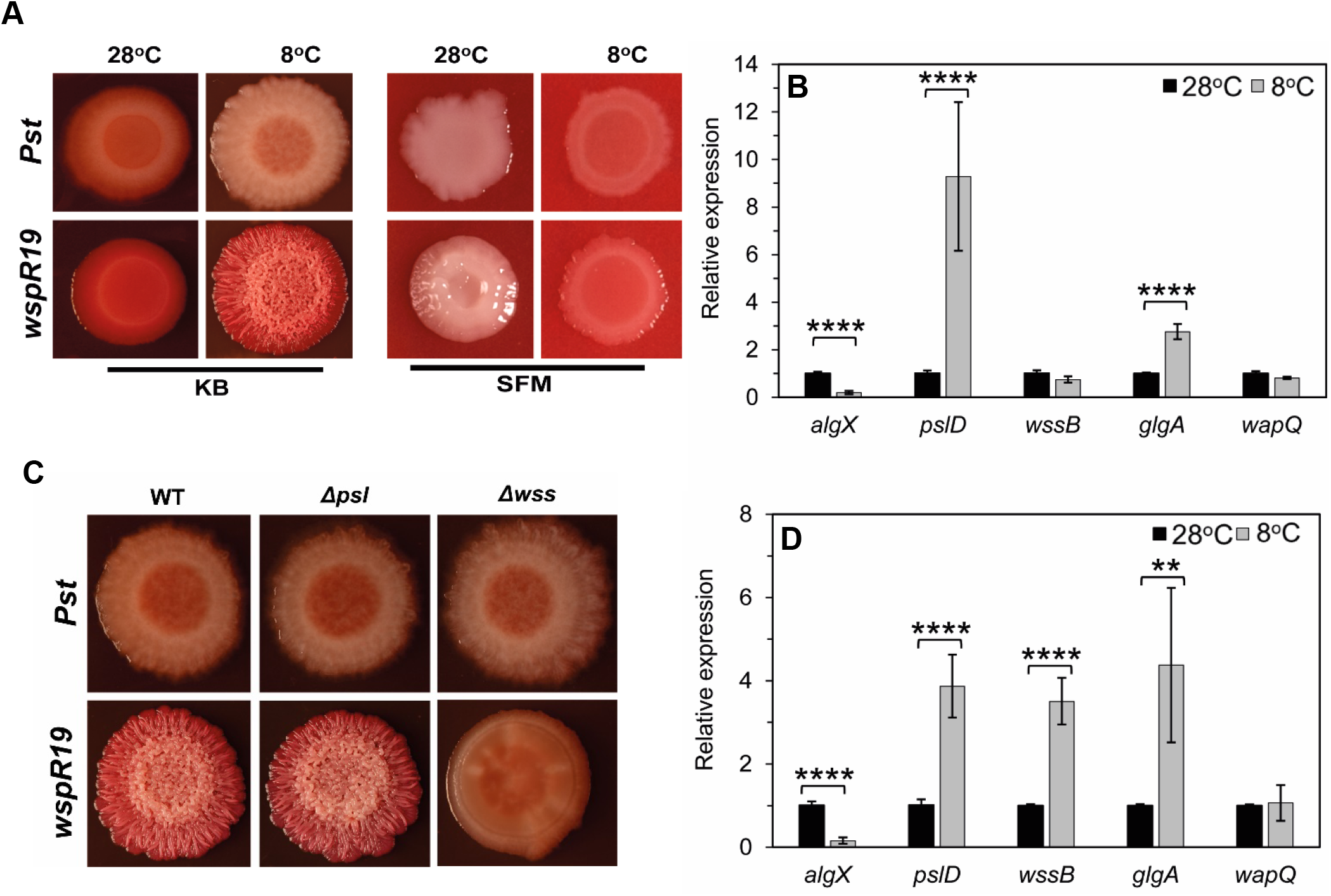
Wss, Psl and α-glucan production are stimulated by low temperature. (A) Effect of temperature on *Pst* colony morphology on CR supplemented media. WT *Pst* and *Pst wspR19* grown on KB and SFM at 28 °C and 8 °C. (B) qRT-PCR analysis of polysaccharide genes in WT *Pst* grown on KB agar at 28 °C (black) and 8 °C (grey). (C) Colony morphology of WT, Δ*psl* and Δ*wss* on KB agar after 5 days of cold treatment at 8 °C. Strains with high cdG (*wspR19*) are shown. (D) qRT-PCR analysis of polysaccharide genes in *Pst wspR19* grown on KB agar at 28 °C (black) and 8 °C (grey). Values represented are mean ± SD with n= 2 (three technical replicates each). Significance was determined by student’s t-test ** indicates p<0.005 and **** indicates p<0.0001). (A) & (C): Similar results were obtained in two independent experiments and a representative image is shown.

The wrinkled colony phenotype on KB was dependent on Wss alone (Fig. 3C). Conversely, growth at 8 °C saw an almost complete abolition of the alginate-dependent, wrinkled mucoid phenotype on SFM plates (Fig. 3A & S9A&B). This was further supported by the decreased levels of *algX* mRNA seen at low temperature (Fig. 3B & 3D). The Wss-dependent wrinkled morphology did not manifest on SFM plates at 8 °C, suggesting that Wss production is nutrient dependent in *Pst* in a similar manner to Psl and alginate (Fig. 3A & S9B).

### α-glucan and alginate production are stimulated by global carbon-storage regulators and surface association

The *treS & treY/Z* operons have previously been implicated in *Pst* leaf surface survival (Freeman *et al*., 2010), suggesting that their expression is likely to be most relevant during epiphytic growth. To investigate this further, we first determined the abundance of key metabolites in *Pst* by ^1^H-NMR spectroscopy. WT *Pst* grown in M9 medium accumulated trehalose and maltose 1 phosphate (M1P) to 0.13 ± 0.03 and 0.30 ± 0.03% of cellular dry weight, respectively (Table 1), alongside a broad NMR peak at 5.41 ppm corresponding to α-glucan (Fig. 4A). Deletion of the *treS* operon alone (*glgE, treS/pep2 & glgB*) not only blocked the production of M1P and α-glucan, as expected, but also led to less trehalose being detected (0.04 ± 0.01%) compared with WT. Deletion of *glgA, treZ, malQ, treY* and *glgX* (Δ*treY/Z*) resulted in no detectable metabolites from these pathways as expected (Fig. 4A and Table 1), whether the other operon was deleted or not.

**Table 1:**
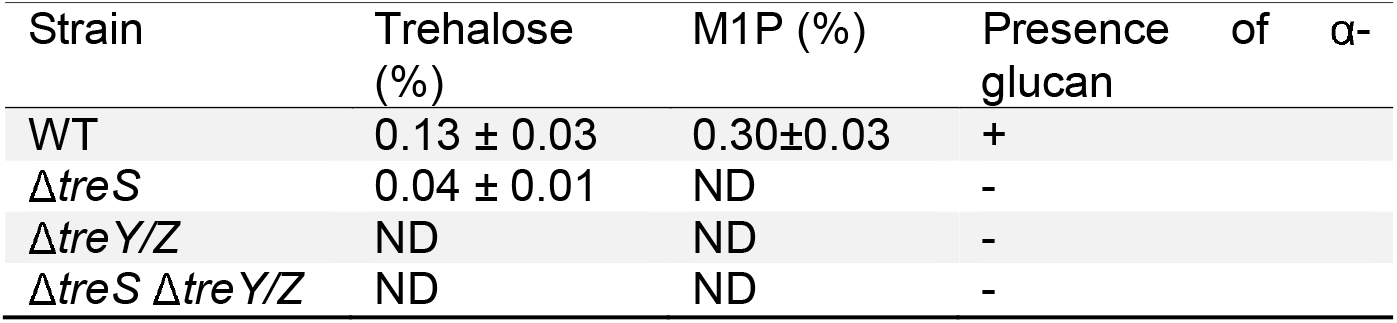
Concentrations of trehalose and M1P produced by *Pst* strains. Metabolites are presented as percentages of cellular dry weight ± standard error (n ≥ 2). ND indicates no measurable metabolite. Presence or absence of α-glucan as indicated by + or – respectively.

| Strain | Trehalose (%) | M1P (%) | Presence of α-glucan |
| --- | --- | --- | --- |
| WT | $0.13 \pm 0.03$ | $0.30 \pm 0.03$ | + |
| $\Delta treS$ | $0.04 \pm 0.01$ | ND | - |
| $\Delta treY/Z$ | ND | ND | - |
| $\Delta treS \Delta treY/Z$ | ND | ND | - |

**Fig. 4.**
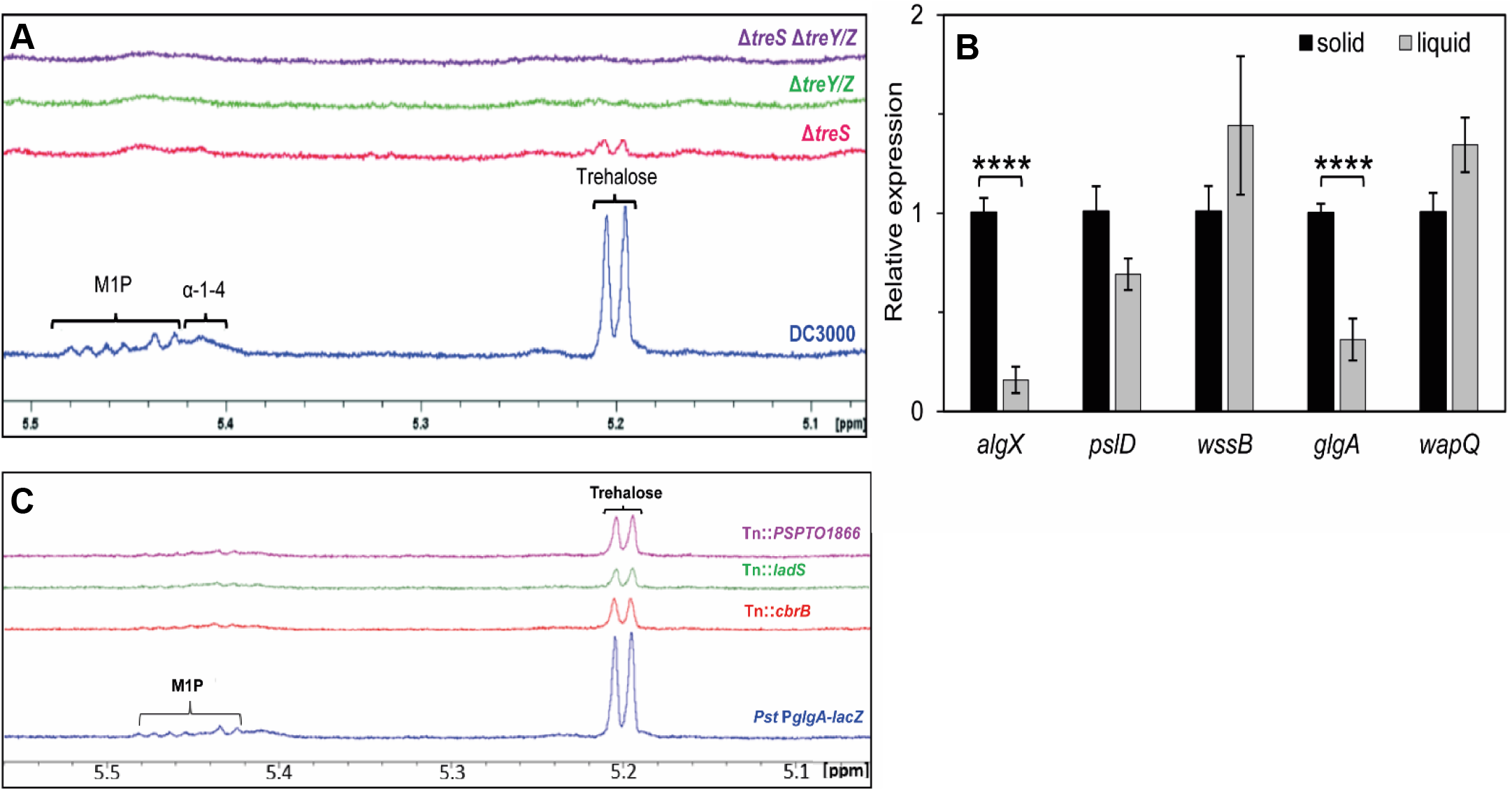
α-glucan and alginate biosynthesis are stimulated by global carbon-storage regulators and surface association. (A) ^1^H-NMR spectra for *Pst* WT (blue) Δ*treS* (pink), Δ*treY/Z* (green) and a double mutant Δ*treS* Δ*treY/Z* (purple). Peaks corresponding to key metabolites are labelled: maltose 1 phosphate (M1P), α-glucan (α-1,4), α-1,1-probable terminal linkage of maltooligosyltrehalose (trehalose). (B) qRT-PCR analysis of polysaccharide genes in WT *Pst* grown on KB agar (solid) vs KB liquid (liquid). Values represented are mean ± SD with n=2 (three technical replicates each). Significance was determined by student’s t-test **** indicates p<0.0001. (C) ^1^H-NMR spectra for the *Pst* P*glgA-lacZ* reporter strain and selected *glgA* regulatory Tn mutants. Peaks corresponding to key metabolites are labelled as in (A).

Next, we used qRT-PCR to measure mRNA abundance for the different polysaccharides genes for *Pst* cells grown on a solid agar surface and in liquid media. Normalized to the *gyrA* internal control, the mRNA level for the α-glucan synthase gene *glgA* was significantly higher in surface-grown compared to liquid-grown *Pst* cells (Fig. 4B). Interestingly, the same pattern of increased mRNA abundance in surface-grown cells was also seen for *algX*, while no differences were seen for the other tested genes.

To further investigate the regulation of *Pst* trehalose/α-glucan gene expression, the 500-bp region upstream of the *glgA* start codon was cloned upstream of *lacZ* and incorporated into the *Pst* chromosome at the neutral *att*::Tn*7* site. The resulting strain (*Pst* P*glgA*-*lacZ*) produced blue colonies on XGal + IPTG plates and was then used to conduct a transposon mutagenesis screen for regulators of *glgA* transcription. Once false positives/negatives and likely indirect hits had been discarded, we identified promising transposon insertions in three regulatory loci: the response regulator *cbrB*, the histidine kinase gene *ladS* and a predicted TetR family transcriptional regulator, *PSPTO1866*. Transposon insertion led to light blue colonies in each case, suggesting that these regulators may function as activators of *glgA* expression.

To test the effects of gene disruption on trehalose and α-glucan biosynthesis, the *Pst* water soluble metabolome was analysed for the three transposon mutants. No significant difference was observed between the levels of analysed metabolites in the soluble metabolomes of WT *Pst* and *Pst* P*glgA*-*lacZ*. Tn::*PSPTO1866* showed a small, but not statistically significant (*p* = 0.06) reduction in trehalose and no change in M1P abundance compared to *Pst* P*glgA*-*lacZ*. Conversely, the metabolomes of Tn::*cbrB* and Tn::*ladS* showed significant decreases in both trehalose and M1P levels (Fig. 4C, Table 2), supporting roles for the global carbon utilisation and chronic/acute lifestyle regulatory proteins LadS (and by extension the Gac/Rsm pathway) and CbrB in stimulating trehalose and α-glucan gene expression (Grenga *et al*., 2017; Sonnleitner *et al*., 2009).

**Table 2:**
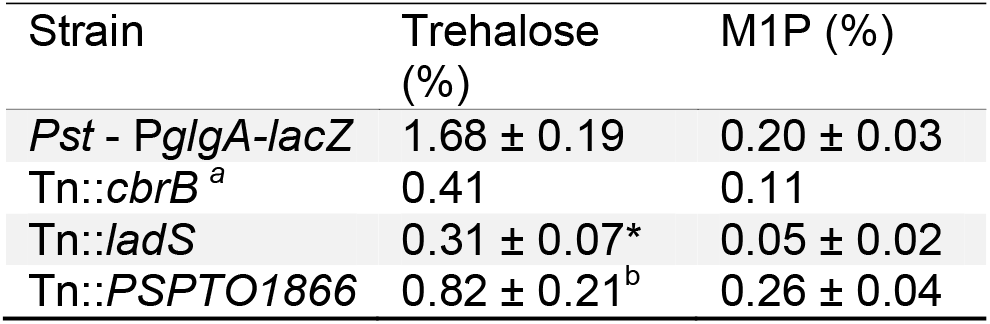
Concentrations of trehalose and M1P produced by *Pst* P*glgA-lacZ* reporter strain and mutants generated by transposon random mutagenesis. Metabolites are presented as percentages of cellular dry weight ± standard error (n ≥ 2), ^a^ indicates n=1. Statistical significance compared to parent strain, * indicates p<0.05 and ^b^ indicates p > 0.05 determined by student’s t-test.

| Strain | Trehalose (%) | M1P (%) |
| --- | --- | --- |
| *Pst* - *PglgA-lacZ* | $1.68 \pm 0.19$ | $0.20 \pm 0.03$ |
| Tn::*cbrB*<sup>a</sup> | 0.41 | 0.11 |
| Tn::*ladS* | $0.31 \pm 0.07^*$ | $0.05 \pm 0.02$ |
| Tn::*PSPTO1866* | $0.82 \pm 0.21^b$ | $0.26 \pm 0.04$ |

**Table 3.** List of strains used in this study

| **Strain name** | **Description** | **Reference** |
| --- | --- | --- |
| WT | Rifampicin resistant *Ps*t considered as WT | (Buell *et al.*, 2003) |
| $\Delta alg$ | Strain with *algG* (PSPTO1238) and *algX* (PSPTO1237) deletion | (Lammertz *et al.*, 2019) |
| $\Delta psl$ | Strain with *pslD* (PSPTO3531) and *pslE* (PSPTO3532) deletion | (Lammertz *et al.*, 2019) |
| $\Delta wss$ | Strain with *wssB* (PSPTO1027) and *wssC* (PSPTO1028) deletion | (Lammertz *et al.*, 2019) |
| $\Delta treS$ | Strain with *glgE* (PSPTO2760), *treS* (PSPTO2761) and *glgB* (PSPTO2762) deletion | This study |
| $\Delta treY/Z$ | Strain with *glgA* (PSPTO3125), *treZ* (PSPTO3126) *malQ* (PSPTO3127), *treY* (PSPTO3128 and (PSPTO3129) and *glgX* (PSPTO3130) deletion | This study |
| $\Delta treS \Delta treY/Z$ | A combination of $\Delta treS$ and $\Delta treY/Z$ | This study |
| $\Delta alg \Delta psl$ | Strain with *algG-X* (PSPTO1238, PSPTO1237) and *pslD-E* (PSPTO3531, PSPTO3532) deletion | (Lammertz *et al.*, 2019) |
| $\Delta alg \Delta wss$ | Strain with *algG* (PSPTO1238) and *algX* (PSPTO1237) and *wssB-C* (PSPTO1027, PSPTO1028) deletion | (Lammertz *et al.*, 2019) |
| $\Delta psl \Delta wss$ | Strain with *pslD-E* (PSPTO3531, PSPTO3532) and *wssB-C* (PSPTO1027, PSPTO1028) deletion | (Lammertz *et al.*, 2019) |
| $\Delta alg \Delta treS \Delta treY/Z$ | Strain with *algG-X* (PSPTO1238, PSPTO1237) deletion combined with $\Delta treS \Delta treY/Z$ | (Lammertz *et al.*, 2019) |
| $\Delta EPS (\Delta alg \Delta psl \Delta wss)$ | Strain with *algG-X*, *pslD-E* and *wssB-C* deletion | (Lammertz *et al.*, 2019) |
| $\Delta wapQ$ ($\Delta PSPTO4998$) | Strain with deletion of an ortholog of *wapQ* ($\Delta PSPTO4998$) | This study |
| $\Delta EPS + \Delta wapQ$ | A combination of $\Delta EPS$ and $\Delta wapQ$ | This study |
| *Ps*t PglgA-lacZ | Reporter strain developed by genome integration of the construct into neutral site using mini Tn7. Reporter construct was generated by cloning lacZ gene under the control of *glgA* promoter. | This study |
| Tn::*cbrB* | *Ps*t PglgA-lacZ strain with identified transposon insertion in *cbrB* gene | This study |
| Tn::*ladS* | *Ps*t PglgA-lacZ strain with identified transposon insertion in *ladS* gene | This study |
| Tn::*PSPTO1866* | *Ps*t PglgA-lacZ strain with identified transposon insertion in *PSPTO1866* gene | This study |
| **Plasmids** |  |  |
| **pTS1** | Tet<sup>R</sup>, suicide vector; *ColE1*-replicon, *IncP-1*, *Mob*, *lacZ* | (Scott *et al.*, 2017) |
| **pBBR-wspR19** | Kan<sup>R</sup>, resistant and di-guanylate cyclase (*wspR19*) expressing plasmid | (Pfeilmeier *et al.*, 2016b) |

**Table 4.**
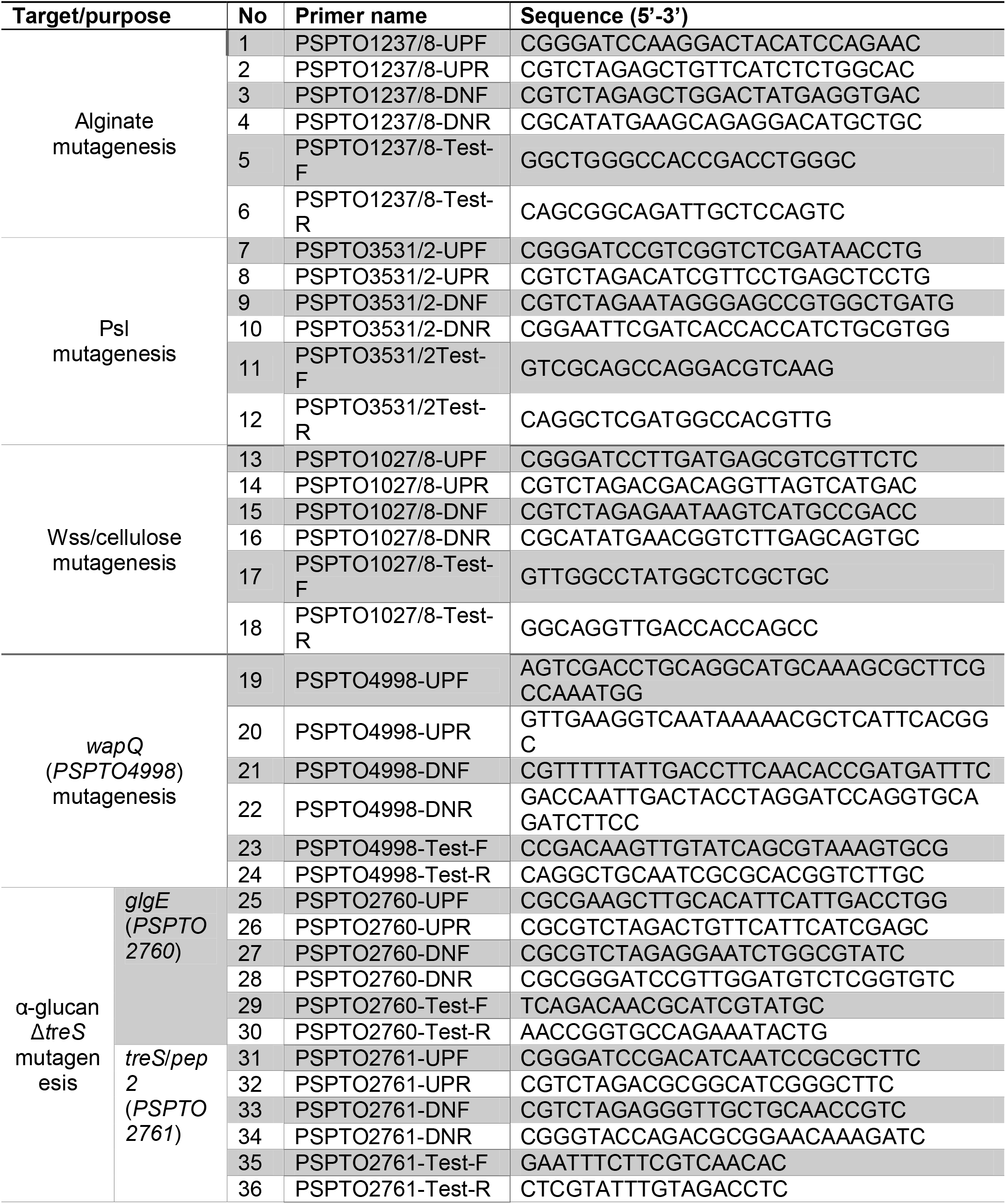
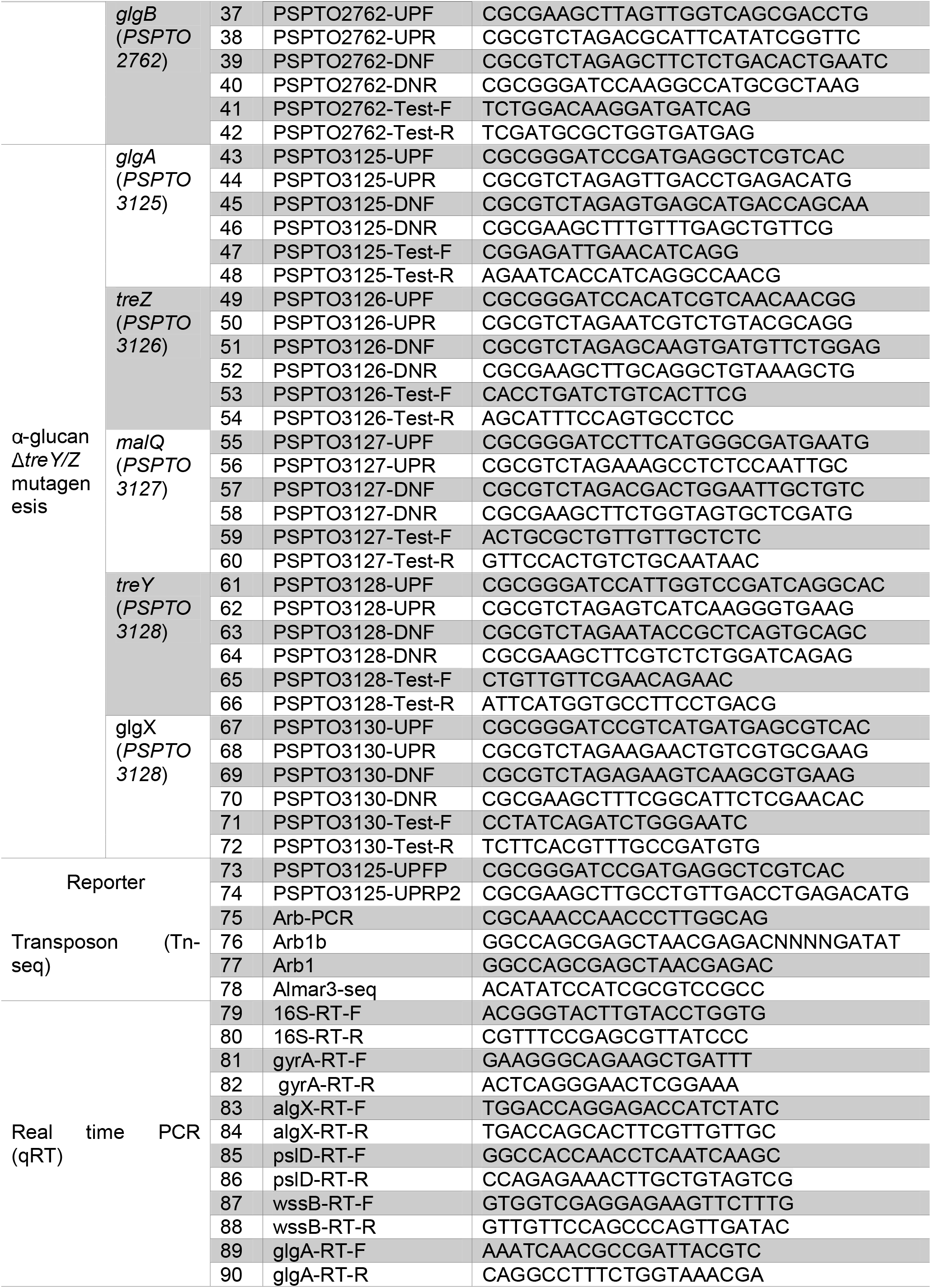
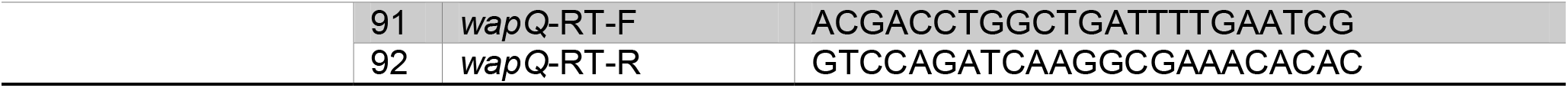
List of primers used in this study

### Disruption of trehalose, α-glucan, and alginate production leads to stress sensitivity and compromised plant infection

To investigate the roles of trehalose and α-glucan in *Pst* during osmotic stress, WT *Pst* and Δ*treS* Δ*treY/Z* were grown in M9 in the presence and absence of 0.35 M NaCl. While both strains exhibited a longer lag-phase and attenuated growth rate when cultured in 0.35 M NaCl, the Δ*treS* Δ*treY/Z* mutant showed substantially increased osmotic sensitivity relative to WT, as expected with the production of trehalose being blocked (Fig. 5A). Next, we analysed the relative desiccation stress tolerance of Δ*treS* Δ*treY/Z* and WT *Pst* as described previously (Woodcock *et al*., 2021). CFU recovered after exposure to 100% and 75% RH were counted, analysed using linear mixed modelling and represented as predicted means of log_10_(CFU ml^-1^). The sensitivity of a strain to reduced humidity was calculated as the difference between its mean log_10_(CFU ml^-1^) at 100% and at 75% RH. A greater response to lower RH of a mutant compared to WT translates to a more desiccation-sensitive strain. Following incubation of bacterial spots on filter discs at 100% RH, 5.83 log_10_(CFU ml^-1^) WT cells were recovered against 5.23 log_10_(CFU ml^-1^) at 75% RH, equating to a desiccation response of approximately 0.60 log_10_(CFU ml-1) (Fig. 5B). The equivalent values for Δ*treS* Δ*treY/Z* were 5.92 (100% RH) and 4.47 (75% RH) log_10_(CFU ml^-1^), giving a desiccation response of approximately 1.50 log_10_(CFU ml^-1^) (Fig. 5B). Based on our recent analysis of PA01 (Woodcock *et al*., 2021), it is likely that this desiccation-sensitive phenotype is linked to the loss of α-glucan production.

**Fig. 5.**
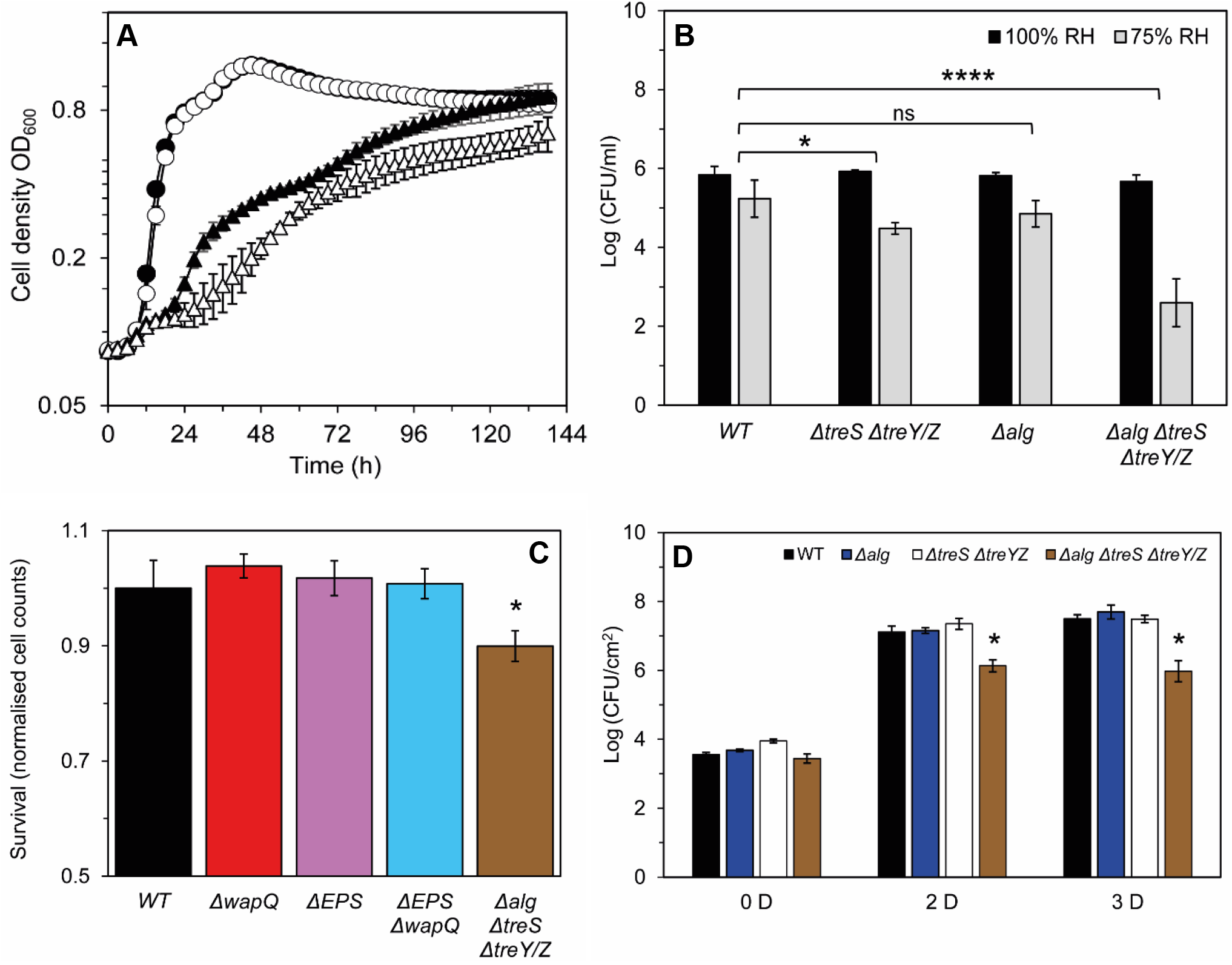
Disruption of trehalose, α-glucan, and alginate production leads to stress sensitivity and compromised plant infection. (A) Growth of WT and Δ*treS* Δ*treY/Z* in M9 medium (circles), WT (solid), Δ*treS* Δ*treY/Z* (open) and in M9 medium supplemented with 0.35 M NaCl (triangles), WT (solid), Δ*treS* Δ*treY/Z* (open). Similar results were obtained for three biologically independent experiments. Data shown is mean ± SD of three technical replicates. (B) Desiccation stress survival assays for strains exposed to 100% RH (black) and 75% (grey) for 2 hours. Data shown is mean ± SD of three biological replicates. (C) Surface survival assay on tomato leaf discs after 24 h. Recovered CFU are normalized against average recovered *Pst* WT CFU for selected mutants. (D) Bacterial load following spray infection at different days post infection [D]. Bars represent the following strains: WT *Pst* (black), Δ*treS* Δ*treY/Z* (white), Δ*alg* (blue) and Δ*treS* Δ*treY/Z* Δ*alg* (brown). Similar results were obtained in three biologically independent experiments and a representative is shown. Error bars represent mean ± SD of 3 technical replicates. In each case, significance was determined by student’s t-test, * indicates p<0.05 and **** indicates p<0.0001.

Alginate has also previously been associated with bacterial stress tolerance and epiphytic fitness (Helmann *et al*., 2019; Yu *et al*., 1999). Based on the apparent link between their colony morphology phenotypes on SFM plates and co-expression on solid surface association, we investigated the role of alginate in protecting *Pst* during desiccation. As the Δ*alg* strain yielded a desiccation response of 0.96 log_10_(CFU ml^-1^), which was not significantly different to the response of wild-type *Pst* when analysed using linear mixed modelling (Fig. 5B), we tested whether α-glucan and alginate may exhibit functional redundancy in *Pst*. An alginate + α-glucan mutant (Δ*alg* Δ*treS* Δ*treY/Z*) was generated and exposed to desiccation stress. This strain yielded a highly significant (p ≤ 0.0001) desiccation response of 3.1 log_10_(CFU ml^-1^) (Fig. 5B). Furthermore, the Δ*alg* Δ*treS* Δ*treY/Z* strain showed reduced epiphytic survival on tomato leaf discs, while no significant difference was observed for other tested strains after 24 h (Fig. 5C). This suggests that alginate and α-glucan functionally complement each other during desiccation stress response on plant leaves.

Next, we investigated whether the interaction of trehalose/α-glucan and alginate is also relevant during plant infection. Col-0 plants were spray infected with the different Δ*alg* and Δ*treS* Δ*treY/Z* mutants monitored over three days’ growth without watering. In agreement with our previous assay (Fig. 1A), a lack of neither alginate nor trehalose/α-glucan alone affected the course of infection. However, we saw a significant decrease in bacterial cell counts for infections with Δ*alg* Δ*treS* Δ*treY/Z* both 2-and 3-days post-infection (Fig. 5D). Together, our results suggest that alginate and α-glucan work together in *Pst* to mediate both desiccation stress tolerance and plant infectivity.

### Disrupting both LPS and EPS alters *Pst* leaf adhesion and compromises plant infection

Several studies have suggested links between the LPS and EPS pathways in *Pseudomonas* spp. (McDonald *et al*., 2009). Given the lack of impact of single gene deletions on Col-0 infection and the complementarity observed between alginate and α-glucan, we tested the effect of disrupting both LPS and EPS biosynthesis on leaf surface interaction and plant infection. Following initial trial assays, Arabidopsis leaves proved too small for reliable measurements of surface association, so tomato leaves (cultivar Moneymaker) were used to examine *Pst* adhesion to and survival on leaf surfaces. In leaf-disc adhesion assays (Arrebola *et al*., 2015), the Δ*EPS* Δ*wapQ* mutant showed a statistically significant reduction (P<0.05) in adhesion to tomato leaf discs compared to the other tested strains (Fig. 6A). Consistent with this, the Δ*EPS* Δ*wapQ* mutant displayed visibly compromised disease phenotypes on Col-0 plants, alongside significantly lower bacterial counts relative to WT *Pst*, the ΔEPS triple mutant or Δ*wapQ* after 3 and 5 days (Fig. 6B). This suggests that the altered surface attachment phenotype seen with this mutant translates into compromised plant infections.

**Fig. 6.**
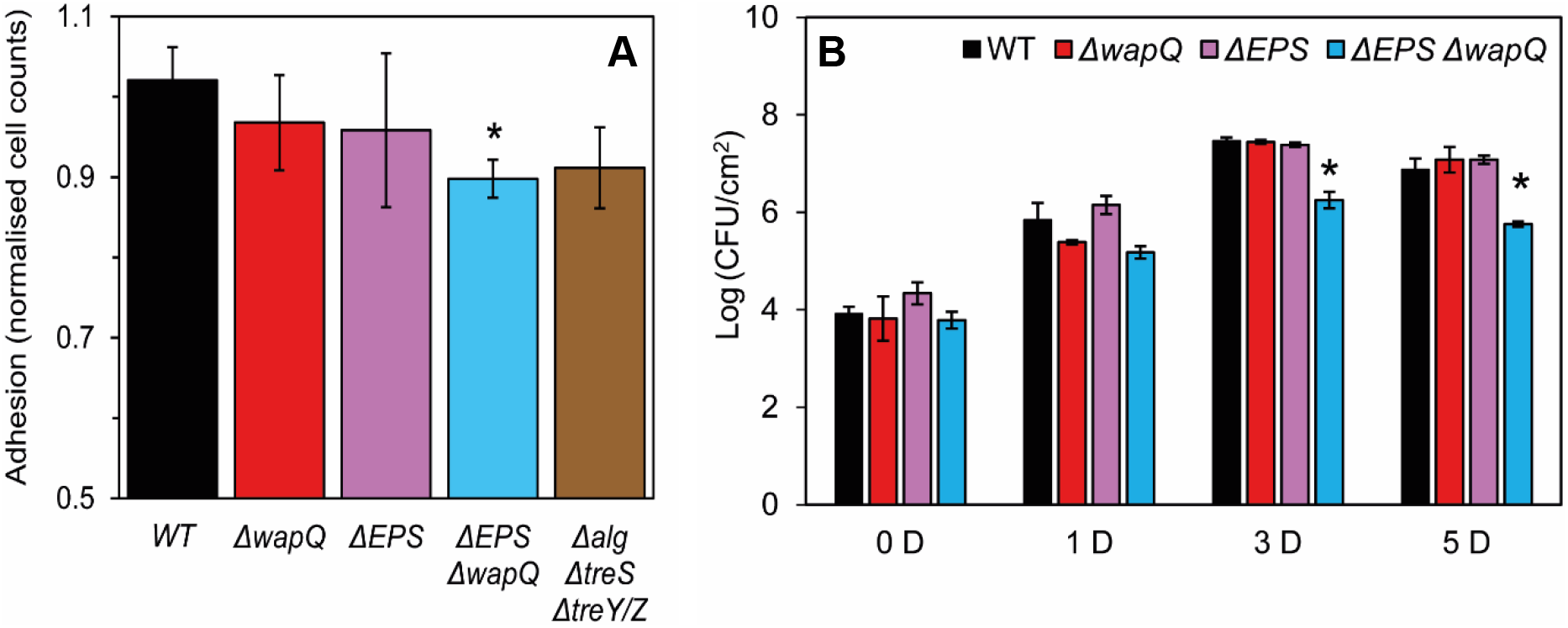
Disrupting both LPS and EPS alters *Pst* leaf adhesion and compromises plant infection. (A) Adhesion to tomato leaf surfaces. Graph denotes number of cells adhered to leaf surfaces after 5h normalized to average number of adhered WT *Pst* cells. Error bars represent mean ± SD of 3 technical replicates. (B) Bacterial load following spray infection at different days post infection [D]. Bars represent the following strains: WT *Pst* (black), Δ*wapQ* (red), Δ*EPS* (purple) and Δ*EPS* Δ*wapQ* (blue). Similar results were obtained in three biologically independent experiments and a representative is shown. Error bars represent mean ± SD of three technical replicates. In each case, the * represents *p*<0.05 significance level determined by student’s t-test.

## Discussion

Phyllosphere colonizing bacterial pathogens such as *P. syringae* face an array of environmental challenges on external plant surfaces (Djonovic *et al*., 2013; Freeman *et al*., 2013; Pfeilmeier *et al*., 2016b; Wu *et al*., 2012). Phytopathogenic bacteria must be able to tolerate rapid shifts in humidity and osmotic pressure, to adhere to leaf surfaces under mechanical disturbance, and to respond effectively to changes in temperature and nutrient availability, among other cues. Systems that facilitate bacterial survival and persistence in the face of these challenges thus play important roles in enabling infection (Pfeilmeier *et al*., 2016a). EPS pathways and cell envelope lipopolysaccharides are ubiquitous in *P. syringae* genomes (Winsor *et al*., 2016) and are among the first bacterial molecules to interact with the host plant (King and Roberts, 2016; Mhedbi-Hajri *et al*., 2011). Polysaccharide biosynthesis has been linked to epiphytic survival, stress tolerance and biofilm formation in various phytopathogenic bacteria (Pfeilmeier *et al*., 2016a). Furthermore, as major components of the biofilm matrix, polysaccharides play important roles not just in leaf surface survival but also during the transition to more acute stages of pathogenicity (Dunger *et al*., 2007; Pfeilmeier *et al*., 2016b; Xin *et al*., 2018; Yu *et al*., 1999).

In this study, we examine the regulation of and interactions between five major polysaccharide pathways in *Pst* and determine their importance to plant infection and epiphytic survival. While EPS & LPS pathways have been implicated in phytopathogen virulence, stress response and biofilm formation (Pfeilmeier *et al*., 2016a), previous studies have generally focussed on the impact of individual polysaccharides (Arrebola *et al*., 2015; Dunger *et al*., 2007; Heredia-Ponce *et al*., 2020). The importance of interactions between the different polysaccharide pathways for plant infection and epiphytic survival is currently poorly understood. Our results suggest that polysaccharide production is tightly regulated and coordinated in *Pst*, with deployment of each polysaccharide system dependent on a number of shared environmental cues (Figure 7). We observed a substantial degree of regulatory and functional redundancy: each external input stimulated multiple polysaccharide pathways, and infection/survival phenotypes were only observed for *Pst* mutants missing several coordinating pathways.

**Fig. 7.**
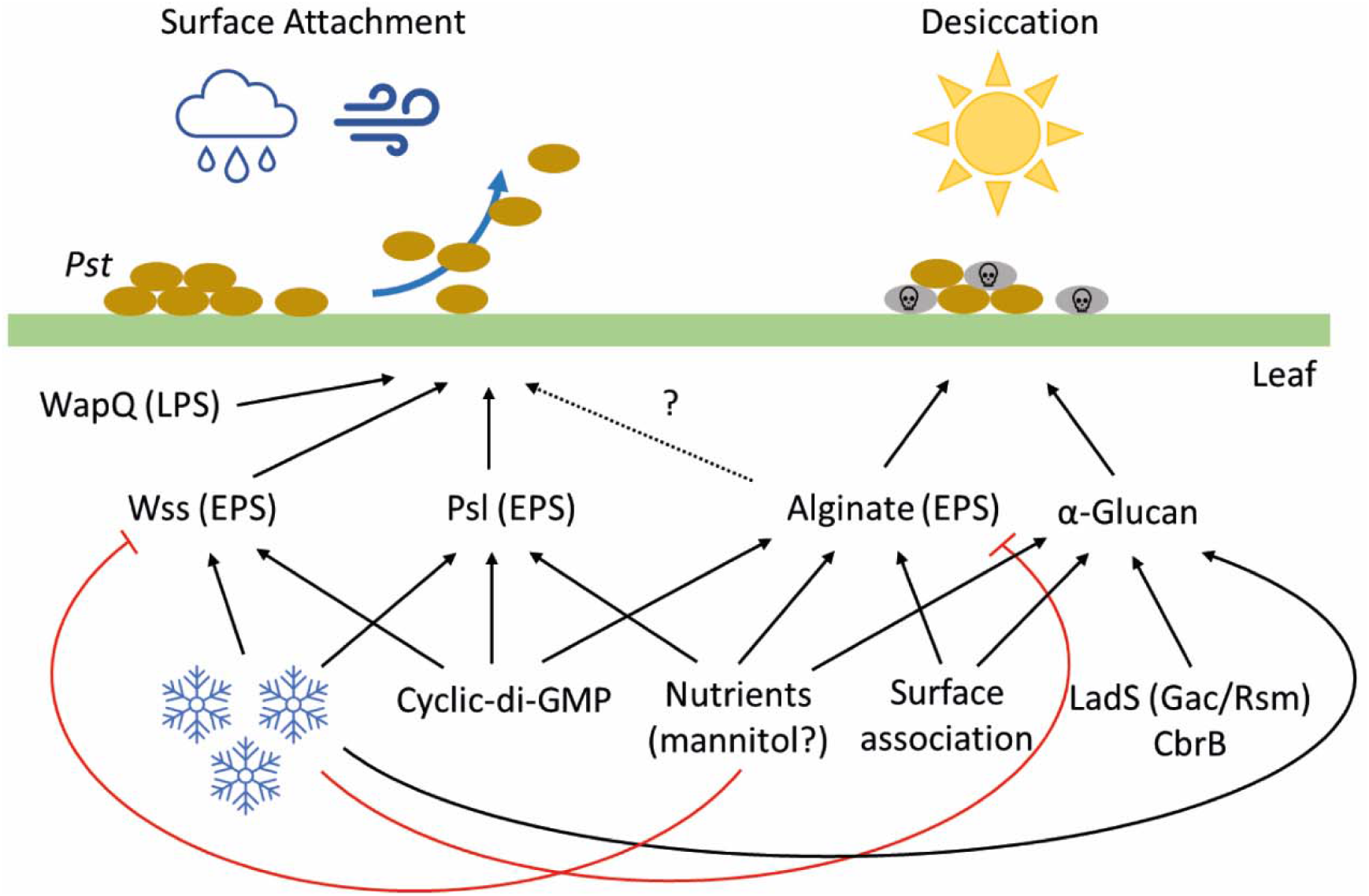
A model for polysaccharide deployment during plant infection by *P. syringae Pst*. Positive or stimulatory interactions are denoted by black arrows. Negative/suppressive interactions are shown by red arrows. An uncertain interaction is indicated with a dashed arrow. Blue snowflakes denote low temperature, brown ovals indicate *Pst* cells and grey ovals indicate dying cells.

CdG conditionally stimulates production of all three *Pst* EPS molecules. However, unlike in other *Pseudomonas* spp. where increased cdG levels lead to constitutive EPS production and wrinkled, aggregative colony morphologies (Malone *et al*., 2010; Malone *et al*., 2007), cdG overproduction in *Pst* does not lead to major changes in colony morphology under standard laboratory conditions. Rather, *Pst* EPS production is highly dependent on external environmental cues, with production of the two major structural EPS molecules; Wss and alginate, stimulated by low temperature and changes to nutrient availability respectively. In each case, the formation of Wss or alginate-driven wrinkled colonies requires both increased cdG production and the correct growth environment. The two phenotypes are mutually exclusive, with Wss linked to growth on KB at low temperatures and alginate to SFM media and surface association, but with little/no effect seen for gene deletions in the other condition.

Psl, which maintains the architecture of mucoid, wrinkly colonies on SFM plates, is stimulated both by low temperature and a switch to SFM growth media, suggesting that Psl is linked to both the alginate and Wss-mediated lifestyles. A central role for Psl in mediating the *Pst* transition from an epiphytic, biofilm lifestyle to more acute pathogenesis was supported by the apparent increase in disease symptoms seen for the Δ*psl* mutant during Col-0 leaf infections. Similar observations have been reported for a *Salmonella enterica* cellulose mutant, and upon Psl or cellulose disruption in *P. syringae* UMAF0158 (Heredia-Ponce *et al*., 2020; Pontes *et al*., 2015). The mechanism by which Psl suppresses disease symptoms during infection, and the relevance of this for bacterial fitness in the plant environment remains to be determined.

Expression of *glgA*; the first gene of the *treY/Z* operon in *Pst*, is stimulated by surface association, SFM media and the global regulatory proteins CbrB and LadS. The response regulator CbrB and its cognate signal kinase CbrA control carbon metabolism, biofilm formation and stress tolerance (Amador *et al*., 2010; Nishijyo *et al*., 2001) by inducing expression of RpoN-dependent genes such as the sRNA *crcZ* (García-Mauriño *et al*., 2013), which acts to antagonise the carbon catabolite repressor Crc (Moreno *et al*., 2009). The Gac/Rsm system controls processes including biofilm, motility, virulence, and the stress response at the level of mRNA translation (Brencic *et al*., 2009; Chambers and Sauer, 2013). In *Pseudomonas* spp., the Gac/Rsm system is controlled by a series of accessory sensor kinases including LadS, which acts as a positive regulator of GacAS activity (Chambonnier *et al*., 2016).

Trehalose/α-glucan and alginate production appear to be subject to a shared regulatory hierarchy in *Pseudomonas* spp. CbrA/CbrB has been implicated in controlling alginate production in *P. fluorescens* (Ertesvåg *et al*., 2017), and ChIP-seq analysis of the *P. putida* genome identified a CbrB binding site in the *algD* promoter (Barroso *et al*., 2018). Similarly, the Gac/Rsm system regulates alginate production in both *Azotobacter* and *Pseudomonas* (Manzo *et al*., 2011; Romero *et al*., 2018), with the anti-sigma factor gene *mucA* an mRNA target for Rsm proteins in *P. aeruginosa*. CbrA/CbrB and LadS are positive regulators of both alginate and trehalose/α-glucan biosynthesis (Ertesvåg *et al*., 2017; Romero *et al*., 2018), which makes sense in the context of their coordinated roles in protecting the cell from desiccation stress. Further research is needed to determine the extent to which the well-established alginate regulon (Wozniak and Ohman, 1994) also controls trehalose and α-glucan biosynthesis.

Trehalose/α-Glucan, and potentially also alginate individually contribute to *Pst* desiccation stress tolerance (Fig. 5B), although the effects on survival or infectivity of disrupting either pathway alone were minimal. However, we saw evidence for a substantial degree of functional redundancy between the two pathways, with trehalose/α-glucan/alginate triple mutants becoming highly sensitive to desiccation stress under laboratory conditions. This *in vitro* desiccation sensitivity translated to decreased epiphytic surface survival (Fig. 5C) and compromised plant infection (Fig. 5D). We also observed functional redundancy between the WapQ LPS kinase and the *Pst* EPS pathways. Leaf attachment and plant infection were unaffected for Δ*wapQ*, or upon disruption of the *wss, psl* and *alg* operons. However, disruption of all four operons together led to significantly reduced leaf attachment and compromised Col-0 infection (Fig. 6A & B).

The emerging picture from our research is that *Pst* uses the coordinated deployment of distinct sets of polysaccharides to address different environmental challenges (Fig. 7). The first of these;, desiccation and osmostress response, are addressed by the deployment of alginate, trehalose, and α-glucan in response to surface contact and changes to nutrient availability, under the control of the Gac/Rsm and CbrA/B regulatory pathways. Conversely, when attachment to leaf surfaces is a high priority, EPS pathways and particularly Wss are upregulated in response to reduced temperatures and under the control of cdG. These two responses appear to be mutually exclusive, with conditions that stimulate Wss production repressing the production of alginate, and *vice versa*.

Our data also suggest that in the context of these broad regulatory groups, *Pst* fine-tunes the deployment of individual polysaccharides to create an optimal response to the environment. For example, *glg* and *alg* regulation are closely aligned, except at low temperatures, where *alg* mRNA is reduced but *glgA* levels increase. Similarly, Psl appear to contribute to both alginate and Wss biofilm formation and is stimulated by SFM and low temperatures. Finally, cdG signalling appears to stimulate several *Pst* polysaccharide pathways, although transcription-level regulation is only apparent for the *wss* locus. Further research is needed to fully understand the regulatory underpinnings of *Pst* polysaccharide deployment during plant infection.

## Author contributions

PSK wrote the manuscript and conducted genetic manipulation, phenotyping, qRT-PCR analysis, and plant infection assays. SP conducted genetic manipulation and plant infection assays. SDW conducted genetic manipulation, phenotyping, plant infection assays and NMR analysis. CZ and SB conceived and designed the study, secured funding, and supervised the research. JGM conceived and designed the study, secured funding, supervised the research, and wrote the manuscript.

## Acknowledgements

Research in the JGM, CZ and SB labs was supported by UKRI-BBSRC Institute Strategic Program Grants BB/J004553/1 (BIO), BB/J004561/1 (MET) and BBS/E/J/000PR9797 (Plant Health) to the John Innes Centre and The Sainsbury Laboratory. PSK was additionally supported by UKRI-BBSRC Grant BB/T004363/1. SDW was funded by a BBSRC Doctoral Training Partnership (BB/J014524/1) PhD studentship. SP was supported by a Norwich Research Park PhD studentship.

## Abbreviations

*Pst*: *Pseudomonas syringae* pv. *tomato* DC3000
LPS: lipopolysaccharide
EPS: exopolysaccharide
PBS: phosphate buffered saline
KB: King’s B medium
L: Lennox medium with 0.1% glucose
SFM: soy flour mannitol medium
M9: M9 minimal nutrient medium
cdG: cyclic di GMP
WapQ: orthologue of LPS kinase WapG/WaaQ/InaA
Waap: Kdo/Waap family LPS kinase
Waal: LPS biosynthesis protein
WbpL: glycosyltransferase/LPS biosynthesis protein
RPM: revolutions per minute
RH: relative humidity
CFU: colony-forming units
cDNA: complementary DNA
CR: Congo Red dye
X-gal: 5-bromo-4-chloro-3-indolyl β-D-galactopyranoside
ddH_2_O: double distilled H_2_O
Δ: deletion of specified gene(s) or operon
G1P: glucose 1-phosphate
G6P: glucose 6-phosphate
M1P: maltose 1-phosphate
OD_600_: optical density at λ 600 nm
NMR: nuclear magnetic resonance
TMSP: trimethylsilyl propanoic acid
SD: standard deviation
CbrB: member of the CbrAB two component regulatory system.

